# Genetic characteristics of human papillomavirus type 16, 18, 52 and 58 in southern China

**DOI:** 10.1101/2021.04.27.438890

**Authors:** Yuee Zu, Zhihua Ou, Dan Wu, Wei Liu, Liwen Liu, Di Wu, Yanping Zhao, Peidi Ren, Yanqing Zhang, Wangsheng Li, Shujin Fu, Yongchun Wen, Xianchu Cai, Wenbo Liao, Chunyu Geng, Hongcheng Zhou, Xiaman Wang, Haorong Lu, Huanhuan Peng, Na Liu, Shida Zhu, Jiyang Liu, Dongbo Wang, Junhua Li

**Author notes:** These authors contributed equally to this work (Yuee Zu, Zhihua Ou, Dan Wu, Wei Liu, Liwen Liu, Di Wu, Yanping Zhao, Peidi Ren).

## Abstract

Persistent infections of high-risk human papillomaviruses (HPVs) are the leading cause of cervical cancers. We collected cervical exfoliated cell samples from females in Changsha city, Hunan Province and obtained 358 viral genomes of four major HPV types, including HPV 16 (n=82), 18 (n=35), 52 (n=121) and 58 (n=100). The lineage/sublineage distribution of the four HPVs confirmed previous epidemiological reports, with the predominant prevailing sublineage as A4 (50%), A1 (37%) and A3 (13%) for HPV16, A1 (83%) for HPV18, B2 (86%) for HPV52 and A1 (65%), A3 (19%) and A2 (12%) for HPV58. We also identified two potentially novel HPV18 sublineages, i.e. A6 and A7. Virus mutation analysis further revealed the presence of HPV16 and HPV58 strains associated with potentially high oncogenicity. These findings expanded our knowledge on the HPV genetic diversity in China, providing valuable evidence to facilitate HPV DNA screening, vaccine effectiveness evaluation and control strategy development.

## Introduction

Human papillomaviruses (HPVs) are double-stranded DNA viruses responsible for nearly all cervical cancers and related to multiple types of other cancers [1]. Of a big family with over 200 different genotypes identified so far, more than 12 HPV types were carcinogenic, among which HPV16 and HPV18 were responsible for over 70% of the cervical cancer cases [2–4]. Meanwhile, HPV52 and HPV58 were dominantly prevailing in East Asian countries, causing more precancer and invasive cancer in this region than elsewhere [5]. Cervical cancer, as the fourth major cancer in women, affected an estimated 570,000 individuals and caused 311,000 deaths in 2018 [6]. To eliminate cervical cancer as a global public health problem, the World Health Organization has raised several targets to be fulfilled by every country by 2035 [7], including 90% vaccination coverage among girls below 15 years of age, 70% screening coverage in 35-45-year-old women and 90% treatment coverage for precancerous and cancer patients. However, the implementation between high-income and low-income countries is significantly different. While 85% of those girls in high-income countries have been covered by national HPV vaccination programs already, only 20-30% of those in low or medium-income countries are covered. Screening implementation also encounters the same dilemma. Therefore, 90% of the cervical cancer deaths occurred in low- or mid-income countries where the preventive measures are least widely practiced [8,9]. A sharp contrast could be found between the US and China. In 2015, the incidence of cervical cancer was 12,900 with 4,100 deaths in the US, while in China, the numbers were estimated as 98,900 and 30,500 respectively [10,11]. The high incidence and mortality of cervical cancer in China was due to the low coverage in both HPV vaccination and cervical cancer screening, under which coverage, the incidence of cervical cancer in 2100 was projected as three times that of 2015 [12]. To meet the WHO target of cervical cancer elimination, both China and other low- and medium-income countries have a long way to go.

The HPV genomes usually encode one long control region (LCR), six open reading frames (ORFs) containing E1, E2, E4, E5, E6 and E7, and the late ORFs expressing L1 and L2 capsid proteins [13]. To better distinguish viral heterogeneity, researchers have designated HPV of the same type into lineage and sublineages, which requires complete genomic nucleotide differences of 1%-10% and 0.5%-1%, respectively [14]. Four lineages (A, B, C and D) and sixteen sublineages of HPV16 (A1-A4, B1-B4, C1-C4, D1-D4) [14–16], three lineages (A, B and C) and nine sublineages (A1-A5, B1-B3, C) of HPV18 [14], four lineages and seven sublineages (A1-A2, B1-B2, C1-C2, D) of HPV52 [14], and four lineages and eight sublineages (A1-A3, B1-B2, C, D1-D2) of HPV58 have been identified so far [14].

HPV sublineages varied both in geographical regions and carcinogenicity. Based on over 7,000 HPV16 positive samples from 52 countries, it was found that the predominant global sublineage was A1, though some other regional predominant sublineages also existed [15]. In East Asia, the regional prevalent HPV16 sublineages were A3 and A4, and sublineage A3, A4 and lineage D may have higher cancer risks than A1 and A2 [15–17]. In Africa, while HPV16 A1/A2 were still prevalent, lineages B, C and D were also common [15]. HPV18 also displayed geographical heterogeneity, with lineage A dominant around the world except in Africa where lineage B was more popular. Within HPV18 lineage A, sublineage A1 was predominant in East Asian and the Pacific [18]. No distinctive cancer risks were identified among the different lineage/sublineages of HPV18, probably due to the limited sample sizes involved in the investigations [18]. As for HPV52, lineage B was predominant in Asia, while lineage A was the predominant lineage in the other continents [19]. For HPV58, lineage A (especially A2) was predominant globally, but lineage C and D were also common in Africa, accounting for 39.1% and 8.7% of all the samples from Africa [20]. It was also reported that the HPV58 A3 sublineage containing E7:T20I/G63S mutations had higher oncogenicity [21,22]. While full genomes were not always available for lineage/sublineage study, most of the above studies used partial genomes to designate HPV lineages/sublineages.

Multiple epidemiological investigations have shown that HPV52 and HPV58 were among the most common HPVs infecting woman cervix in China, and may cause more infections than HPV16 and HPV18 in some regions [23–25]. The disease burden caused by HPV52 and HPV58 should not be neglected considering the large population size in China, although their carcinogenicity was inferior to that of HPV16 and HPV18 [26]. Currently, most of the investigations on HPV genomes were carried out in North America, only a small amount of them were from China. Moreover, the lineage/sublineage distribution of HPV types was mainly classified by partial genomes or single genes, and the HPV diversity in southern China remained obscure. Herein, using samples from Changsha, Hunan Province, we aimed to reveal the genetic diversity of the HPV16, 18, 52 and 58 with full viral genome sequences. A comprehensive understanding of the viral characteristics of these four types would facilitate the evaluation of the protection potency of related HPV vaccines in China.

## Results

### Classification of HPV16 genomes from Changsha

In this study, we have obtained 82 HPV16 full genomes from Changsha (Table 1, Supplementary Table S1). Although HPV16 was the most common HPV type causing cervical cancer, HPV16 genomes from China were limited. Combined with public data, we constructed Maximum-Likelihood phylogeny based on full viral genomes to understand the lineage distribution of the Changsha strains. Based on phylogeny clustering and distance comparison, the HPV16 from Changsha city were mainly assigned to sublineage A4 (41, 50.00%), A1 (30, 36.59%), and A3 (11, 13.41%) (Table 1, Figure 1). Some of the strains in sublineage A1 formed two new branches neighboring A3 and A4, which were labelled as cluster M and N in the HPV16 phylogeny (Figure 1). Pairwise sequence distance calculation showed that genomes in the two branches differed from the A1 reference genome by less than 0.5%, indicating that the two branches still belonged to sublineage A1. Besides the sequences from China, HPV16 strains from other countries including the United States, Japan and Thailand, were also identified in these two branches (Figure 1, Supplementary Figure S1).

**Table 1:**
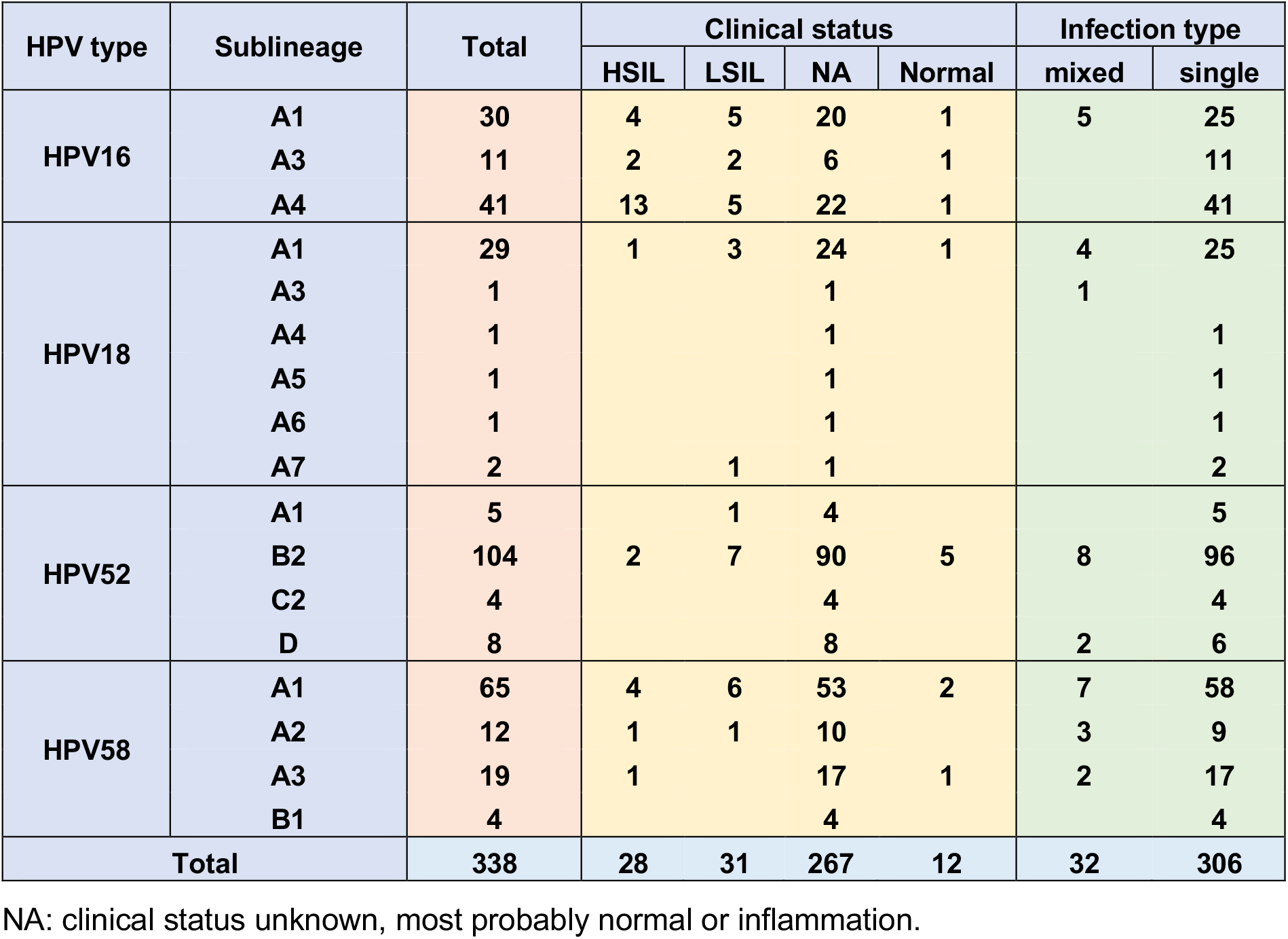
Characteristics of the HPV genomes generated by this study.

**Figure 1:**
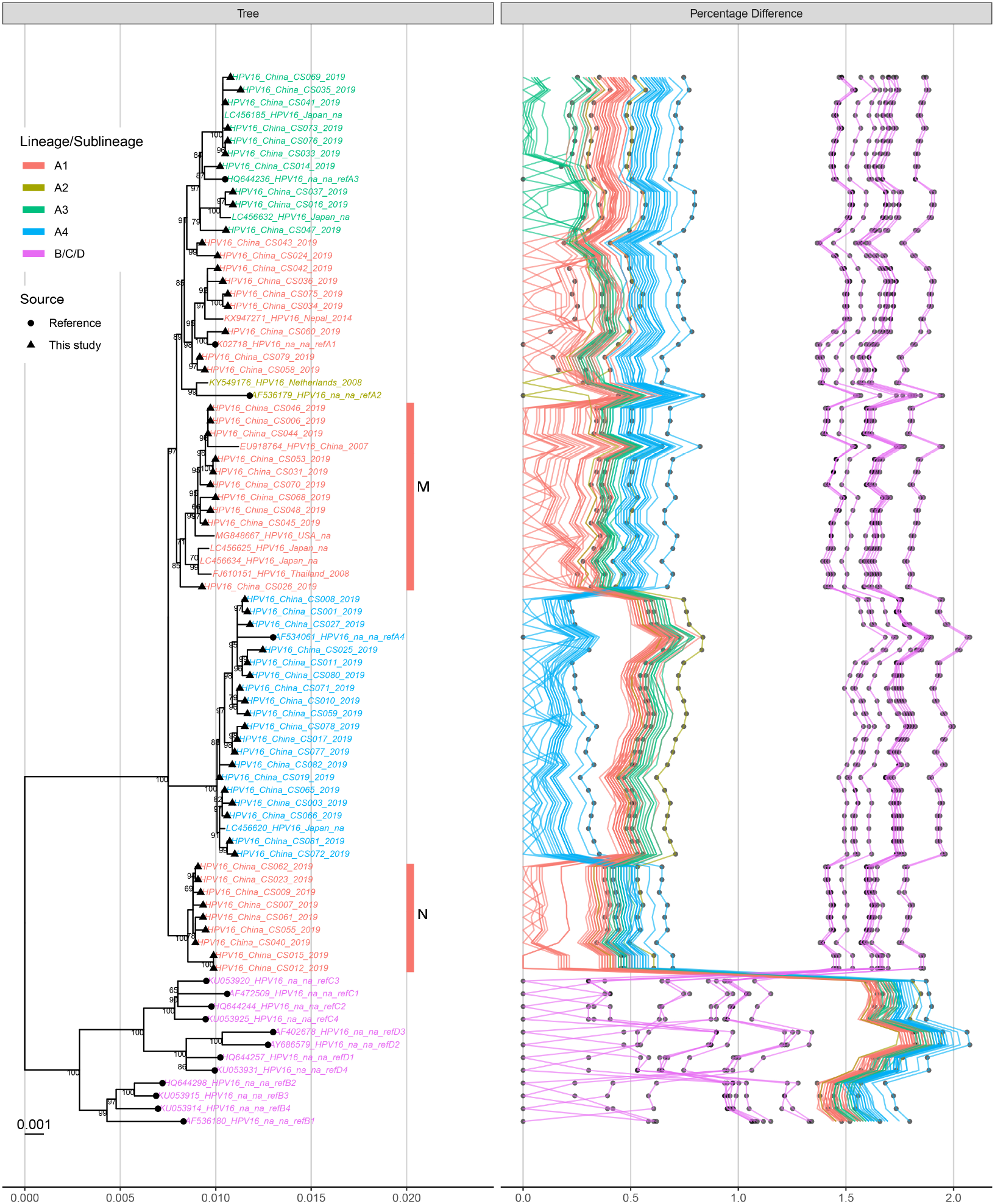
Representative Maximum Likelihood phylogeny of HPV16. HPV genomes generated by this study were combined with those from public database to construct a Maximum Likelihood phylogeny with 1,000 bootstrap tests. Both the phylogenetic tree (left panel) and the pairwise sequence distance (right panel) are shown. Sources of the sequences are indicated by shapes while lineages or sublineages are distinguished by different colors. Bootstrap values over 70 are labeled on nodes.

### Classification of HPV18 genomes from Changsha

For HPV18, we have obtained 35 full genomes from samples in Changsha (Table 1, Supplementary Table S1). Based on phylogeny topology and nucleotide difference, the HPV18 from Changsha city were mainly assigned to sublineage A1 (29, 82.86%). We also identified one strain each for sublineages A3, A4 and A5. Moreover, there were three strains that belonged to lineage A but not assigned to any knonw sublineages (Table 1, Supplementary Table S1, Supplementary Figure S2), which we designated as sublineages A6 and A7 (Figure 2). Sublineage A6 contained three strains, with two from China and one from the Netherlands. These strains had less than 0.5% distance from HPV18 sublineage A3 reference genome but failed to form a monophyletic branch with sublineage A3. Meanwhile, their nucleotide differences from sublineages A1, A2 and A4 were approximately 0.5%. Therefore, we decided to classify this cluster as a new sublineage. Sublineage A7 was a neighboring branch to HPV18 sublineage A5, which contained two strains from Changsha with sequence distances of 0.5%-1% from all the A sublineages. Considering only limited strains were present in sublineage A6 and A7, more epidemiological evidence may be needed to support the presence of the two new HPV18 sublineages.

**Figure 2:**
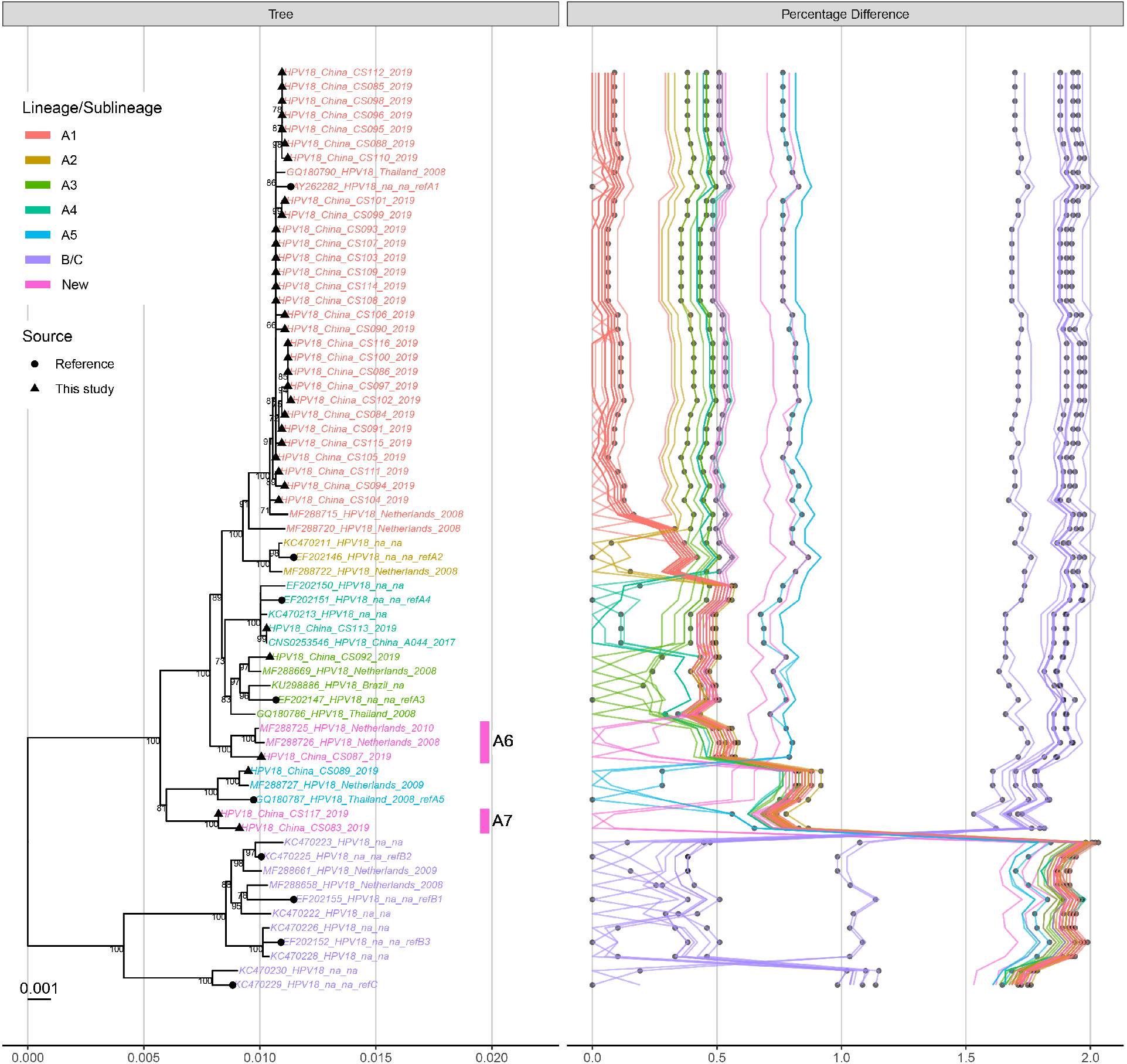
Representative Maximum Likelihood phylogeny of HPV18. Figure legends are the same as Figure 1.

### Classification of HPV52 genomes from Changsha

For HPV52, we have obtained 121 full genomes from samples in Changsha (Table 1, Supplementary Table S1). The HPV52 strains were mainly assigned to sublineage B2 (104, 85.96%, Figure 3, Supplementary Figure S3). Several strains from sublineage A1 (5, 4.13%), C2 (4, 3.31%) and D1 (8, 6.61%) were also detected, implicating the sporadic circulation of multiple lineages in this region. It was intriguing to have HPV52 sublineage D1 detected, which was rarely reported by other studies in Asia.

**Figure 3:**
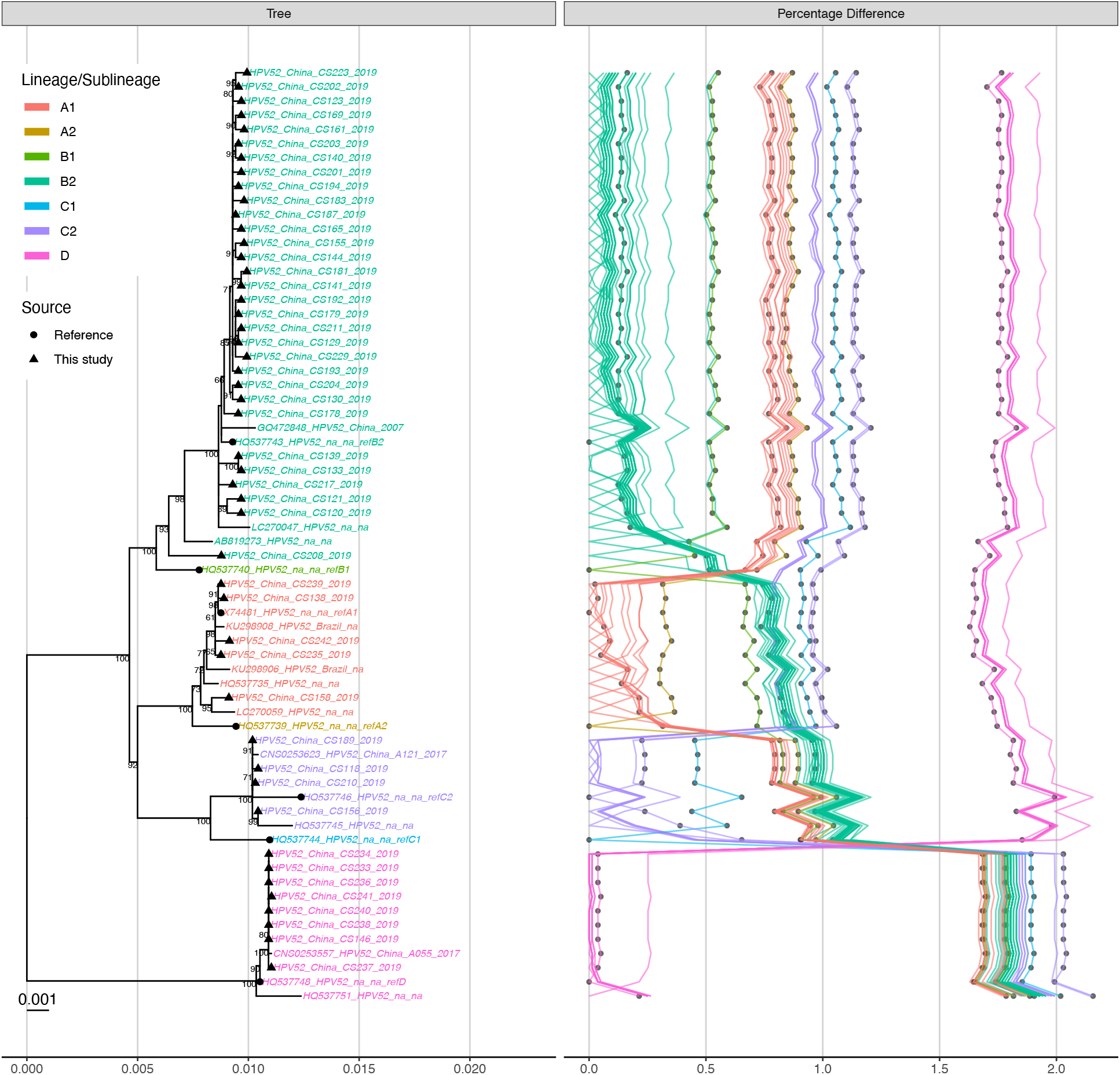
Representative Maximum Likelihood phylogeny of HPV52. Figure legends are the same as Figure 1.

### Classification of HPV58 genomes from Changsha

For HPV58, we obtained 100 full genomes from the Changsha samples (Table 1, Supplementary Table S1). The HPV58 from Changsha city were mainly assigned to sublineage A1 (65, 65.00%), A2 (12, 12.00%) and A3 (19, 19.00%) (Figure 4, Supplementary Figure S4). Four strains belonged to sublineage B1 were also detected.

**Figure 4:**
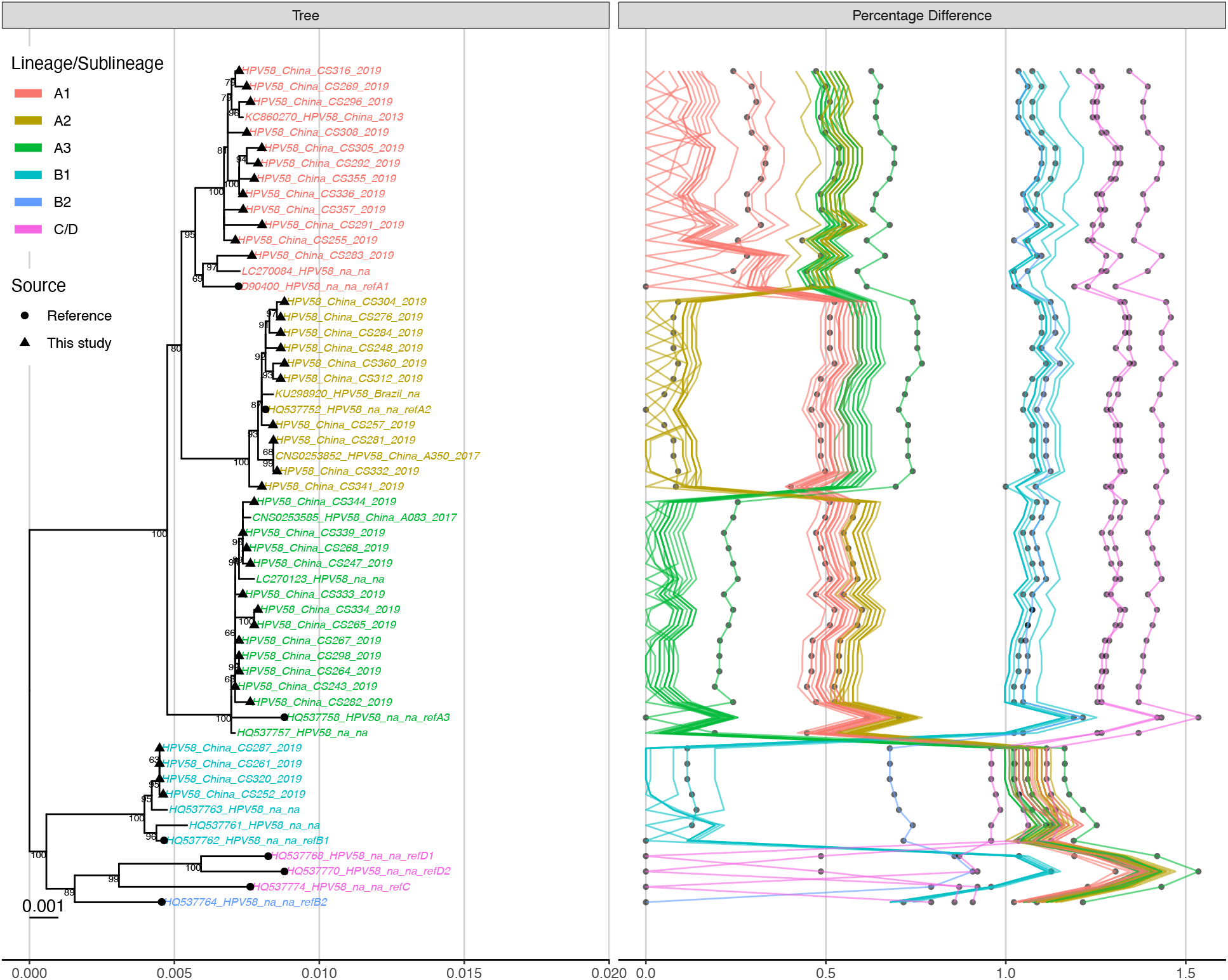
Representative Maximum Likelihood phylogeny of HPV58. Figurelegends are the same as Figure 1.

### Lineage/sublineage conservative mutations identified for HPV16, 18, 52 and 58

Lots of the epidemiological studies on the lineage or sublineage distributions of HPVs were relied on partial genome sequencing and comparison on nucleotide polymorphism. To help refine fixed mutations of lineages or sublineages, we combined the Changsha data with publicly available HPV genomes to conduct genome-wide mutation analysis for the four high-risk HPV types (Supplementary Table S2). Herein, we defined the mutations that occurred in over 98% of the strains belonging to at least one lineage or sublineage as conservative mutations. Mutations in E4 were not shown in the main text because this gene located within E2, but the related details could be found in the supplementary materials.

For HPV16, a total of 210 positions with conservative mutations were identified based on 2,480 full genomes (Table 2, Supplementary Table S3), these included 79 missense and 91 synonymous mutations occurred in seven genes (E1, E2, E5, E6, E7, L1 and L2), and 40 point mutations occurred in the long control region (LCR) and noncoding regions (NCR). Among them, 39 mutations were probably unique to 11 sublineages (Table 3). No unique mutations were identified for sublineages A2, A3 and B1. L1, L2 and LCR contained the most abundant unique mutations, and could be used to distinguish 7, 6 and 6 HPV16 sublineages, respectively. The combination of the unique sites from L1, L2 and LCR would be able to distinguish at least 11 sublineages.

**Table 2:** Distribution of the genomic positions with conservative mutations for HPV16, 18, 52 and 58 lineages or sublineages. Blank cells indicate no detection.

| HPV Type | Mutation Type | E1 | E2 | E5 | E6 | E7 | L1 | L2 | LCR | NCR | Subtotal | Total |
| --- | --- | --- | --- | --- | --- | --- | --- | --- | --- | --- | --- | --- |
| HPV16 | missense mutation | 15 | 25 | 4 | 8 | 1 | 13 | 13 |  |  | 79 | 210 |
|  | synonymous mutation | 32 | 10 | 4 | 2 | 3 | 20 | 20 |  |  | 91 |  |
|  | point mutation |  |  |  |  |  |  |  | 33 | 7 | 40 |  |
| HPV18 | missense mutation | 10 | 21 | 6 | 1 | 3 | 13 | 22 |  |  | 76 | 269 |
|  | synonymous mutation | 34 | 16 | 5 | 9 | 2 | 25 | 34 |  |  | 125 |  |
|  | point mutation |  |  |  |  |  |  |  | 31 | 6 | 37 |  |
|  | deletion |  | 6 |  |  |  |  |  | 7 | 18 | 31 |  |
| HPV52 | missense mutation | 12 | 21 | 7 | 4 | 8 | 9 | 5 |  |  | 66 | 289 |
|  | synonymous mutation | 25 | 12 | 2 | 7 | 3 | 36 | 22 |  |  | 107 |  |
|  | point mutation |  | 1 |  |  |  |  |  | 37 | 7 | 45 |  |
|  | deletion |  |  |  |  |  |  |  | 19 | 21 | 40 |  |
|  | insertion |  |  |  |  |  | 3 |  |  | 28 | 31 |  |
| HPV58 | missense mutation | 12 | 3 |  | 5 | 6 | 17 | 7 |  |  | 50 | 186 |
|  | synonymous mutation | 16 | 8 | 6 | 2 | 4 | 22 | 23 |  |  | 81 |  |
|  | point mutation |  |  |  |  |  |  |  | 37 | 6 | 43 |  |
|  | deletion |  |  |  |  |  |  |  | 12 |  | 12 |  |

**Table 3:**
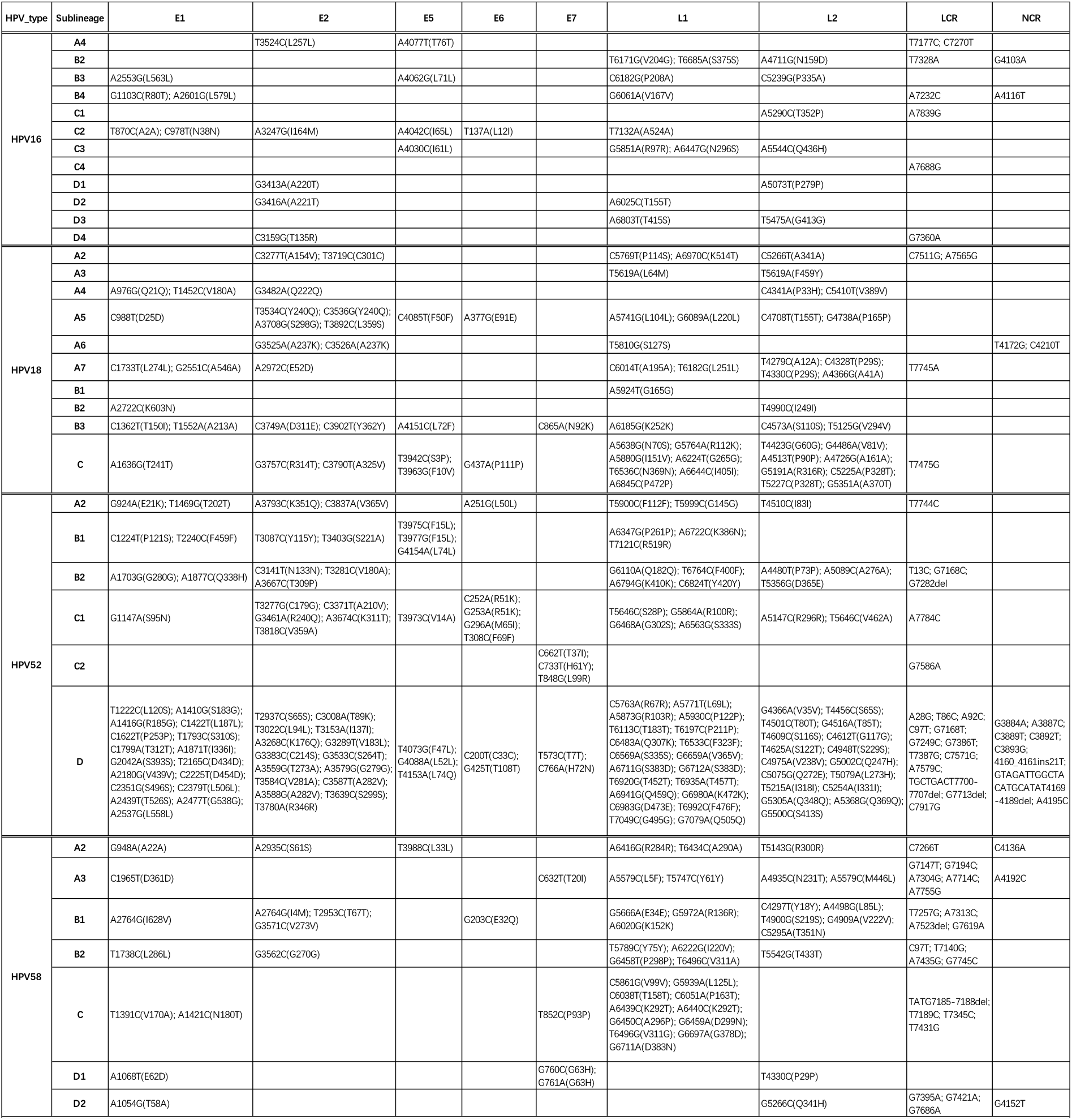
Unique mutations detected in certain lineages/sublineages of HPV 16, 18, 52 and 58.

For HPV18, a total of 269 positions with conservative mutations occurring in at least one lineage or sublineage were identified based on 182 full genomes (Table 2, Supplementary Table S4). These mutations included 76 missense mutations, 125 synonymous mutations and 6 deletions in the coding genes, 37 point mutations and 25 deleted sites occurred in the NCR and LCR (The deletion at one nucleotide position was counted as one deletion site. For example, a 6bp consecutive deletion was counted as 6 deletion sites). A 6bp deletion in the overlapping region of E2, CCTACA3630-3635del (E2:PT272-273del), and A 7bp deletion in LCR (TGTTGTA7234-7240del) was observed in sublineages A5, A7, B1, B2 and B3 and lineage C (Supplementary Table S4). In NCR, an 18bp deletion (3915-3932) occurred in strains of sublineages B1 and B2, and a 7bp deletion (3919-3925) was observed in lineage C only. A total of 74 lineage/sublineage unique mutations were identified (Table 3). L1 and L2 contained the most abundant unique positions, and their combination could distinguish all the HPV18 lineages/sublineages. For the two novel HPV18 sublineages, i.e., A6 and A7, five and ten unique mutations were identified respectively. Because some sublineages of HPV18 contained very few genomes, such as sublineages A5, A6, A7 and lineage C (Supplementary Table S2), it’s possible that some random mutations might be included. Still, for some major sublineages, the unique mutations would be informative, such as T5619A(L1:L64M) for sublineage A3, T1452C(E1:V180A) and C4341A(L2:P33H) for sublineage A4, A5924T(L1:G165G) for sublineage B1, etc.

For HPV52, a total of 289 positions with conservative mutations were identified based on 314 full genomes (Table 2, Supplementary Table S5). These mutations included 66 missense mutations and 107 synonymous mutations in the coding genes, 45 point mutations and 40 deleted sites occurred in the NCR and LCR, and 31 inserted nucleotides in L1 and NCR. Interestingly, a 21bp deletion (4169-4189) in NCR and an 8bp deletion in LCR (7700-7707) was only observed in lineage D (Table 3, Supplementary Table S5). We also found a long insertion comprised of 28 nucleotides between the 4160^th^ and 4161^st^ positions of HPV52 (based on the A1 reference genome), which occurred in all strains of lineage B, C and D. Moreover, lineage D had G at the 21^st^ position of this long insertion, while lineage B and C had T (Table 3). A total of 169 unique mutations were identified. E1, E2, L1 and LCR all contained unique mutations for five sublineages, and the combination of unique sites from L1 and LCR would be able to distinguish all the lineages and sublineages of HPV52. It should be noted that lots of the unique sites belonged to lineage D, which was probably due to the long genetic distances between lineage D and the other lineages.

For HPV58, a total of 186 positions with conservative mutations were identified based on 321 full genomes (Table 2, Supplementary Table S2, Supplementary Table S6). These mutations included 50 missense mutations and 81 synonymous mutations in the coding genes, 43 point mutations in NCR and LCR, and 12 deleted sites in LCR. A 7bp consecutive insertion (7164-7170) in LCR was found in sublineages D1 and D2. A total of 79 unique sites was identified for HPV58, and E1 contained the most abundant unique sites for all the lineages and sublineages (Table 3).

## Discussion

Previous studies on HPV sublineages were mainly conducted using partial gene sequences and may fail to reveal the variation profiles of full genomes. In this study, we have conducted genomic surveillance on four high-risk HPV types in Changsha city to explore the genetic diversity of HPV16, 18, 52 and 52 in southern China in a much higher resolution using complete viral genomes. We showed that A4, A1 and A3 were the major HPV16 sublineages circulating in Changsha, similar to our previous findings based on samples from eastern China [27]. Our genomic findings were also consistent with the epidemiological reports by other studies using partial gene sequencing [28,29]. We have also identified two minor clusters within HPV16 sublineage A1. While the two clusters were mainly formed by strains from China, Japan and Thailand (Supplementary Figure S1), the exact distribution of cluster M and N in East Asian countries remains to be clarified. Expanding genomic surveillance in Asia would further reveal the geographical structure of HPV16 variants. Complete genomes of HPV18, 52 and 58 were relatively limited in public database, with 150-250 genomes of each type from Asia, Europe and the Americas available for this investigation [27]. Our study showed high prevalence of HPV18 sublineage A1 in Changsha, China (Table 1), in accordance with findings by Chen et al [18]. We have also detected two potential novel sublineages of HPV18 in this region (Figure 2, Supplementary Figure S2), which was not identified in other regions. Interestingly, our previous genomic study on HPV18 in eastern China showed a Chinese cluster in sublineage A4 [27], which was not found in this study (Supplementary Figure S2). This suggests that the genetic diversity of HPV18 may be more divergent than what has been reported and that different regions in China may have their unique variants. HPV52 displayed higher divergence than HPV16 and HPV18, as the isolates belonged to four different lineages. Based on available genome data, B2 was the most prevalent sublineage for HPV52 both in China and the world (Supplementary Figure S3) [30,31]. Although in low ratios, HPV52 strains of lineage C and D were continuously detected in China [27,32], whether this is due to long-term maintenance or recent population movement remains unknown. HPV58 mainly belonged to lineage A, with A1 as the dominant sublineage (Supplementary Figure S4), consistent with the overall distribution of HPV58 in Asia [20,30,32]. Our phylogenetic classification revealed several potentially high-risk HPV sublineages in Changsha, including HPV16 A4 [16] and HPV58 A3 [33], raising concerns about their circulation status among the Chinese population. Our study provided useful information on the sublineage distribution of four major high-risk HPV types, which would serve as useful resources for the surveillance and control of HPVs in Chinese females.

The sublineage conservative mutations identified in our study also provided evidence on the genetic divergence and potential high-risk markers in the HPV variants in China. For example, the T178G/A (E6: D25E/E) mutation, especially T178G, has been repeatedly detected in prevailing HPV16 variants in many provinces of China, including Hubei, Xinjiang, Liaoning, Harbin, Zhejiang, Yunnan, Taiwan, Hong Kong, etc [29,34–38]. Here, T178G was uniquely detected in 40 out of 41 (97.6%) sublineage A4 strains from Changsha, while T178A was uniquely detected in 7 out of 11 (63.6%) strains of sublineage A3, suggesting that T178G mutation may be a common mutation for sublineage A4. Therefore, it is possible that the variants containing T178G mutations belong to HPV16 sublineage A4 and that this sublineage may be prevalent in different provinces of China. HPV16 T350G (E6:L83V) was another point mutation that’s frequently mentioned in HPV16 variants [17,37,39], which might be a marker for sublineage D3 [35]. While no genomes of sublineage D3 were obtained from Changsha, we did find 3 strains (10%) of sublineage A1 with the T350G mutation. Moreover, this mutation was also found in sublineage A2 and B4 (Supplementary Table S3), indicating that T350G was not specific to assign lineages or sublineages. It was suggested that HPV16 variants with the E6:D25E mutation in the Japanese population and those with E6:L83V mutation in Swedish women were linked with higher carcinogenicity, but the sample sizes of such studies were relatively small [40,41]. For HPV18, mutations related to carcinogenicity were rarely reported. For HPV52, Choi *et al* reported that A379G (E6 K93R) might be linked with higher disease severity and that this mutation was mainly occurred in sublineage B2 [31]. In our study, we showed that A379G mutation occurred in not only sublineage B2, but also in A2 and B1 (Supplementary Table S5). Other mutations previously found to be associated with sublineage C, such as A706G, G707A, T727G, C733T, G742A and T848G [31], were also detected in the HPV52 strains (n=4) of sublineage C2 in Changsha. Moreover, these mutations were also found to be unique to lineage C (Supplementary Table S5). For HPV58, both epidemiological and experimental evidence indicated that linked mutations of C632T (E7:T20I) and G760A (E7:G63S), mostly found in sublineage A3, were positively associated with high oncogenicity [22,33,42]. While C632T uniquely occurred in 100% (n=19) of the Changsha sublineage A3 strains, G760A was found in all strains of sublineage A3 (n=19) and B1 (n=4) (Supplementary Table S6). Unfortunately, our dataset failed to provide supportive evidence regarding the carcinogenicity of HPV sublineages. Zehbe *et al* pointed out that the carcinogenicity of HPV variants may also depend on the host immune system, for example, the individual difference in the major histocompatibility complex [41]. Current study also showed that some of the risk-associated mutations, such as HPV16 E6:L83V, were not lineage- or sublineage-specific. Whether the lineage/sublineage classification or mutation detection better relates with disease progression remains to be explored. Moreover, with the availability of more abundant complete viral genomes, the lineage- or sublineage-specific mutations would also be further refined.

With an aim to understand the genetic diversity of the four major high-risk HPVs, the samples were randomly selected based on their infection types but not strictly taken in accordance with their epidemiological prevalence. Although the infection and clinical information for samples were summarized (Table 1), such data were not eligible for any correlation analysis, which presents as one limitation of this study. In our mutation analysis, the lineages or sublineages containing the fewer stains tended to have more unique mutations. Certain unique mutations identified for the minor lineages/sublineages may be false positive but could be refined by increasing the dataset in the future. The mutation profile might help reveal the genetic divergence of the HPV strains in southern China but some of them might not be common characteristics for strains from other areas.

Due to the low mutation rate of DNA viruses and the transmission route of HPVs, our study in Changsha would help reveal the HPV genomic heterogeneity in Hunan and even southern China. However, for regions with unique ethnic groups and coastal regions with frequent population exchange, the HPV diversity may be more complicated. Moreover, HPVs displayed the highest diversity in Africa [15,20,43], where genomic surveillance was relatively limited. With the initiation of the cervical cancer elimination campaign, continuing HPV screening and gradual application of vaccines are expected around the world. It would be critical to understand the viral genetics before and after vaccine coverage, so as to better evaluate the effectiveness of vaccines, make appropriate adjustments to vaccine combination of HPV types, select the best reference vaccine candidates and improve other related preventive strategies. The HPV genomes generated by our study would serve as valuable baseline reference data for the control of HPVs in women in southern China. The conservative mutations identified may also facilitate large-scale lineage/sublineage classification of HPV16, 18 52 and 58 during epidemiological study. Conducting genomic surveillance in different regions of China and other regions of the world would help us better understand the baseline of viral activity, hence deciding the best practice for screening strategies and vaccine application.

## Methods and Materials

### Sample collection

Exfoliated cervical cell samples were obtained from women aged from 35 to 64 participating in the Cervical Cancer Screening Program from May to June 2019 in Changsha City, Hunan Province, China. HPV types were determined with BGI SeqHPV Kit (BGI-Shenzhen, China). All the participants consented to the donation of their leftover clinical samples for this investigation. The samples that were positive of HPV16, 18, 52 or 58, regardless of single-type or multiple-type infections, were used for DNA extraction.

### HPV DNA enrichment and sequencing

Genomic DNA was extracted with MagPure Buffy Coat DNA Midi KF Kit (Magen, China). Samples with a total DNA amount of over 400ng were further fragmented to around 200–400 bp with a Covaris LE-220 ultrasonicator. These fragments were then used to construct the sequencing libraries following the instruction of the MGIEasy DNA Library Preparation Reagent Kit (MGI, BGI-Shenzhen, China). These purified fragments were end repaired, adaptor-ligated, and amplificated with 8 PCR cycles. The post-PCR products were used to enrich HPV fragments. HPV RNA probes targeting 18 HPV types (HPV6, 11, 16, 18, 31, 33, 35, 39, 45, 51, 52, 56, 58, 59, 66, 68, 69 and 82) were designed by MyGenostics [44] and synthesized by MGI, BGI-Shenzhen. The hybridization process was carried out according to the manufacturer’s instructions (MGI, Shenzhen, China). Libraries were hybridized with HPV probes at 65 °C for 24 hours to capture HPV-specific fragments. The eluted fragments were amplified by 18 cycles of PCR and then used to generate DNA nanoball-based libraries after circularization and rolling circle amplification. Libraries were sequenced as 100bp paired-end reads on the BGISEQ-500 and MGISEQ-T1 sequencing platform (MGI, Shenzhen, China).

### Complete genome assembly

The raw reads were quality-checked and trimmed with fastp [45]. Deduplicated reads were further removed with BBMap (https://sourceforge.net/projects/bbmap/). The clean reads were mapped to HPV reference genomes with the BWA alignment tool [46]. Reads with both ends aligned to HPV reference genomes were extracted and used for *de novo* assembly with NOVOPlasty [47]. The assembled contigs were locally blasted against a reference genome of the same type to identify the relative genomic location. The genomic fragments were then adjusted to have the same genomic coordination as the reference genome. Reference genomes for the four HPV types investigated in this study were: HPV16, K02718; HPV18, AY262282; HPV52, X74481; HPV58, D90400. It should be noted that the HPV16 reference strain K02718 was downloaded from PaVE [4], which was longer than the GenBank version by two nucleotides.

### Phylogeny reconstruction and data visualization

The best-fit nucleotide substitution models were determined by ModelFinder and the Maximum Likelihood (ML) phylogenies were all constructed with 1,000 implementations of ultrafast bootstrap tests with IQ-TREE. To fully reveal the genetic diversity of the new viral genomes generated by this study, we initially constructed maximum phylogenies of the four HPV types combining genomes from NCBI and CNSA for our preliminary analysis. The total sequences and the nucleotide substitution models were: HPV16, n=2584, GTR+F+I+G4; HPV18, n=182, TVM+F+I+G4; HPV52, n=315, GTR+F+I+G4; HPV58, n=323, TVM+F+I+G4. To improve visual clarity, we selected representative strains to construct the final phylogenies: HPV16, n=83, TVM+F+I+G4; HPV18, n=65, K3Pu+F+I; HPV52, n=66, TVM+F+I; HPV58, n=55, TVM+F+I. The pairwise nucleotide differences were calculated with *seqinR* (*http://seqinr.r-forge.r-project.org/)*. Based on genomic differences and phylogenetic topology, lineage and sublineage classification were performed for four HPV types [14]. Visualization of phylogeny and the associated data were carried out with *ggtree* package in R [48].

### Mutation detection

Mutation detection was conducted by comparing the new genome sequences against the A1 reference genome of each HPV type (HPV16, K02718 from PaVE [4], with E6 started from genomic nucleotide position 104, which was different from the GenBank record; HPV18, AY262282; HPV52, X74481; HPV58, D90400). Only genomes with correct reading frames for all the eight coding regions (E1, E2, E4, E5, E6, E7, L1 and L2) were used for mutation detection. To remove random mutations, only those detected in over 98% sequences of any lineage or sublineage were retained. Mutations that uniquely detected in a specific lineage or sublineage were further identified. Sequences with incorrect reading frame or early stop codons in the coding regions were excluded from the mutation analysis. The numbers of genomes used for mutation analysis were as follows: HPV16, n=2480; HPV18, n=182; HPV52, n=314; HPV58, n=321 (Supplementary Table S2).

### Ethical statement

This study was reviewed and approved by the Institutional Review Board of both Changsha Maternal and Child Care Hospital and Beijing Genomics Institute, Shenzhen, China (BGI-R071-1-T1 & BGI-R071-1-T2). All the participants provided written consent for this study.

## Data availability

The data that support the findings of this study have been deposited into CNSA (CNGB Sequence Archive) of CNGBdb under project number CNP0001700 (https://db.cngb.org/cnsa/).

## Author contributions

Junhua Li, Yuee Zu, Zhihua Ou, Jiyang Liu and Dongbo Wu designed and supervised the study. Yuee Zu, Dan Wu, Liwen Liu, Wenbo Liao, Yanqing Zhang, Yongchun Wen, Zhihua Ou, Yanping Zhao, Xiaman Wang, Huanhuan Peng, Na Liu and Shida Zhu coordinated sample collection. Peidi Ren, Wei Liu and Hongchen Zhou performed DNA extraction and library construction. Wangsheng Li, Shujin Fu, Chunyu Geng, Xianchu Cai and Haorong Lu conducted viral genome sequencing. Zhihua Ou, Di Wu and Wei Liu conducted data analysis. Zhihua Ou, Yanping Zhao, Peidi Ren and Wei Liu wrote the manuscript. Junhua Li, Yuee Zu, Dan Wu and Liwen Liu provided critical comments on the manuscript.

## Acknowledgments

We thank China National GeneBank for providing sequencing service for this project. We warmly thank Ms. Jieyao Yu, Ms. Wei Zhou and Mr. Qineng Li for their assistance in viral genome sequencing. The authors also wish to extend their gratitude to Miss Feiyun Ou and Mr. Geer Xi for their inspirational communications.

## Supplementary Materials

**Supplementary Figure S1: Maximum Likelihood phylogeny of HPV16**. HPV genomes generated by this study were combined with those from public database to construct a Maximum Likelihood phylogeny with 1,000 bootstrap tests using IQ-TREE. Number of genomes: 2,584. Nucleotide substitution model: GTR+F+I+G4. Sources of the sequences are indicated by shapes while lineages or sublineages are distinguished by different colors. Bootstrap values over 70 are labeled on nodes.

**Supplementary Figure S2: Maximum Likelihood phylogeny of HPV18**. Number of genomes: 182. Nucleotide substitution model: TVM+F+I+G4. Figure legends are the same as Supplementary Figure S1.

**Supplementary Figure S3: Maximum Likelihood phylogeny of HPV52**. Number of genomes: 315. Nucleotide substitution model: GTR+F+I+G4. Figure legends are the same as Supplementary Figure S1.

**Supplementary Figure S4: Maximum Likelihood phylogeny of HPV58**. Number of genomes: 323. Nucleotide substitution model: TVM+F+I+G4. Figure legends are the same as Supplementary Figure S1.

**Supplementary Table S1: Classification of the HPV16, 18, 52 and 58 genomes obtain from Changsha**.

**Supplementary Table S2. Lineage/sublineage distribution of the genomes used for mutation detection**.

**Supplementary Table S3. Nucleotide mutations detected in HPV16 full genomes**.

**Supplementary Table S4. Nucleotide mutations detected in HPV18 full genomes**.

**Supplementary Table S5. Nucleotide mutations detected in HPV52 full genomes**.

**Supplementary Table S6. Nucleotide mutations detected in HPV58 full genomes**.

