## Supplementary figures and images for "Genetic characteristics of human papillomavirus type 16, 18, 52 and 58 in southern China"

### Supplementary Figure S1

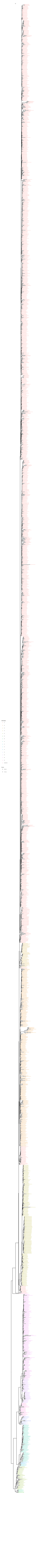

### Supplementary Figure S2

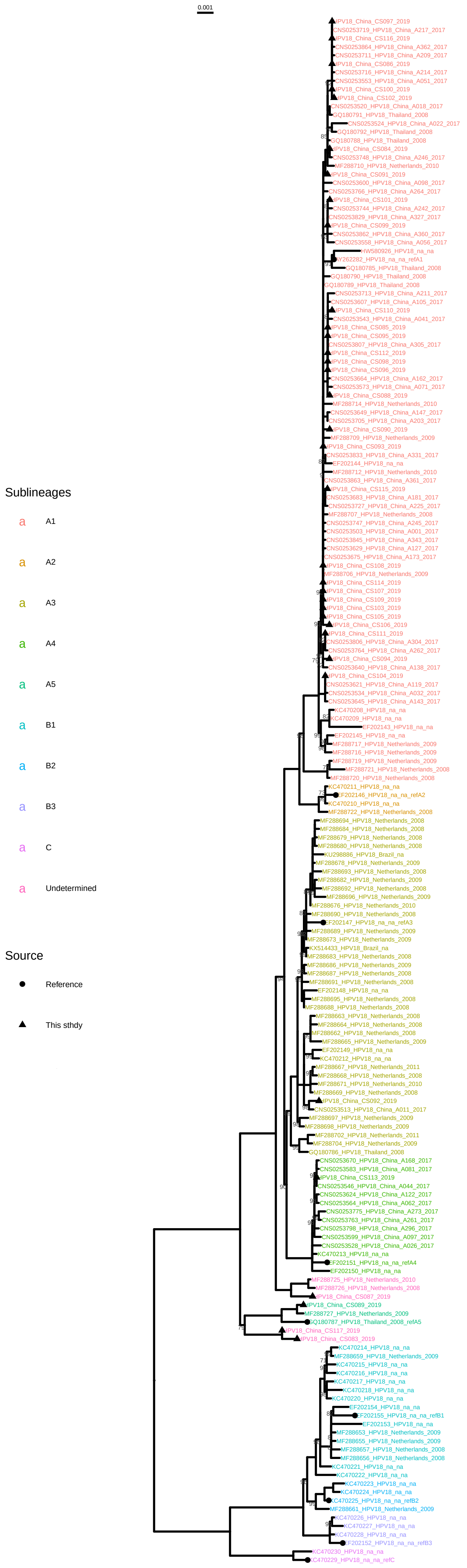

### Supplementary Figure S3

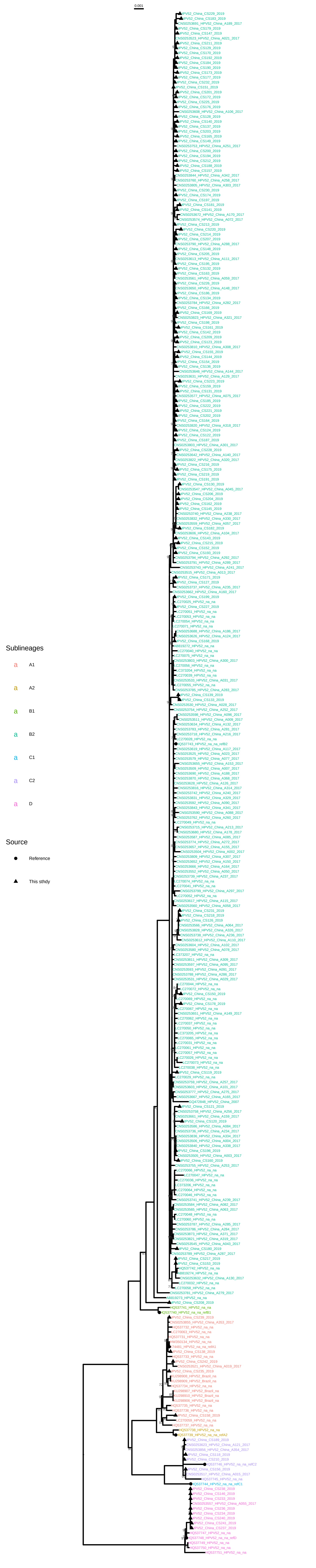

### Supplementary Figure S4

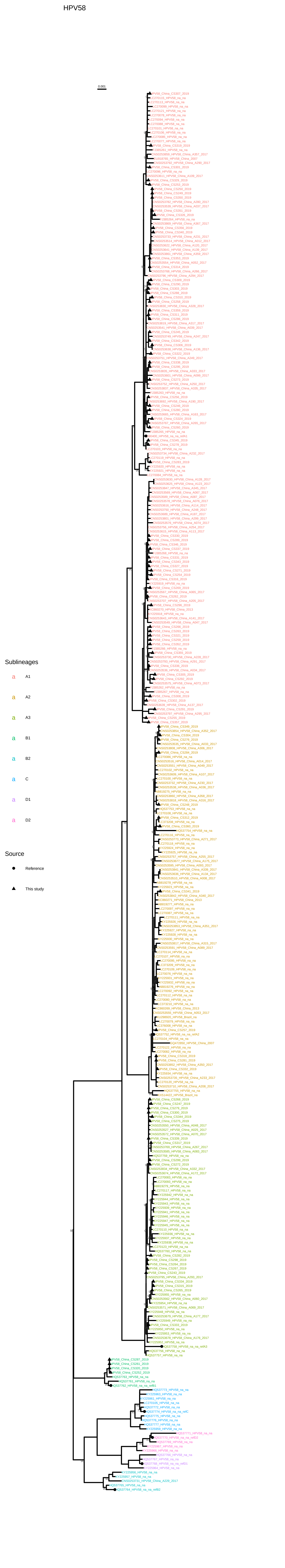
